# Theta-frequency resonance as the scale of input-output mapping in the medial entorhinal cortex

**DOI:** 10.1101/369025

**Authors:** Nupur Katyare, Sujit Sikdar

## Abstract

Grid cell spatial period is thought to be dictated by a mapping between the speed-direction modulated excitatory inputs, and consequent modulation of the firing rate, yet, the exact underlying mechanisms are not known. Here, through experiments on the medial entorhinal cortex stellate cells, subjected to *in-vivo* like stochastic synaptic activity through the dynamic clamp, we show that such mapping can emerge from a theta-frequency resonance in the signal gain, which is HCN sensitive, robust to noise, and is potent enough to modulate the synaptic responses in the theta frequency. This modulation also extends to the corresponding theta-gamma modulation of the firing rate, the slope of whose excitation mediated increase is steeper in the presence of HCN channels. We also show that in the cells devoid of HCN channels, inhibition can emulate their role. Considering the dorso-ventral gradients of HCN and inhibition, which are present aligned to the grid spacing gradient in the medial entorhinal cortex, these findings should be noteworthy.

---

The computational capabilities of Stellate cells of the Medial Entorhinal Cortex (MEC) are evident from their grid-like and border-like spatial representations^1–3^, their necessity for spatial learning^4^, their ability to facilitate contextual representations in the downstream circuits^5^, steeper slopes of phase precession^6^, and their possible contributions to the theta and the gamma rhythms^7^.

Although other excitatory cell types have now been reported to exhibit grid-like firing patterns^1,8^, the strong correlation between the gradients of intrinsic properties and integration time window of stellate cells along the dorso-ventral (DV) axis of the Medial Entorhinal Cortex (MEC)^9–12^, and a similar gradient observed in the spacing of the grid fields^13,14^, is still in need of many mechanistic and theoretical explanations.

Theories explaining grid cell spatial periodicity generally assume a mapping between the speed-direction modulated inputs, and the consequent modulation of the membrane voltage response; accounting for the path integration^12,15–20^. The intrinsic mechanisms in stellate cells conjectured to affect this mapping are mainly the intrinsic theta oscillations^19,21,22^, other HCN mediated mechanisms^9,23,24^ and the membrane resistance and its effects on the time constants^9,12^. Intrinsic theta oscillations have been confirmed in *in-vivo* intracellular recordings of the grid cells^2,3^, and have been reported to be potent enough to influence the spike timings but not to generate the firing fields on their own, which were rather found to be generated by the slow synaptic ramps^2,3^. The involvement of HCN channels is evident from the experiments on the HCN1 knockout mice^25^, which reported expanded grid fields and reduced slope of the theta frequency modulation by speed.

All these observations lead to the following questions: How the ramps and theta oscillations in the membrane potential can emerge from the noisy stream of synaptic inputs? Through which mechanisms can HCN channels influence the mapping between the excitation and the firing rate which is thought to influence the grid spatial periodicity? And what alternate mechanisms operate in the cells devoid of HCN channels?

A way to address these questions is to consider that although the intrinsic theta oscillations can be weakened by the *in-vivo* noise^26^, the underlying HCN dependent resonance mechanisms can still manifest in shaping the cellular responses^27–29^.

To look for such functional resonance mechanisms, we computed signal gain in stellate cells under dynamic clamp mimicked *in-vivo* states, and observed that it indeed displays a strong HCN dependent theta-frequency resonance, which is also robust to the underlying noise. Interestingly, we observed that this resonance in signal gain also reflects in a strong theta modulation of the synaptic noise, and is also accompanied by a faster gamma frequency bursting; which together lead to a steeper theta frequency firing rate modulation by excitation.

While studying this response modulation, we also observed the membrane potential signatures of theta oscillatory activity and the slow synaptic ramps to emerge from an interplay between the HCN channels and the excitatory input kinetics.

We could confirm these results in a computational model of stellate cell subjected to the same input regime, which provided additional insights regarding the role of inhibition as an alternate resonance mechanism in the cellular circuits devoid of HCN channels.

The fact that the dorso-ventral gradients of HCN and inhibition are present in the medial entorhinal cortex^9,11,30^, aligned to the grid spacing gradient^13,14^, further underlines the importance of these results.

Our findings thus indicate that the acuteness and scaling of responses in these cells is critically maintained by the mechanisms of resonance contributed by HCN channels and inhibition.

## RESULTS

### Theta frequency peak in the signal gain

We estimated signal gain of stellate cells when the cells were firing in theta frequency under the mimicked *in-vivo* states. The stellate cells were identified from their electrophysiological responses (**Supplementary Figure 1**) in the whole cell current clamp mode, following which the *in-vivo* states were mimicked through the dynamic clamp^31^ (**Supplementary Figure 2)**. The successful mimicry^31^ was tallied from large voltage fluctuations following a large depolarization (**Fig. 1a**), and ~ 70 % drop in the input resistance (**Fig. 1b**).

**Figure 1.**
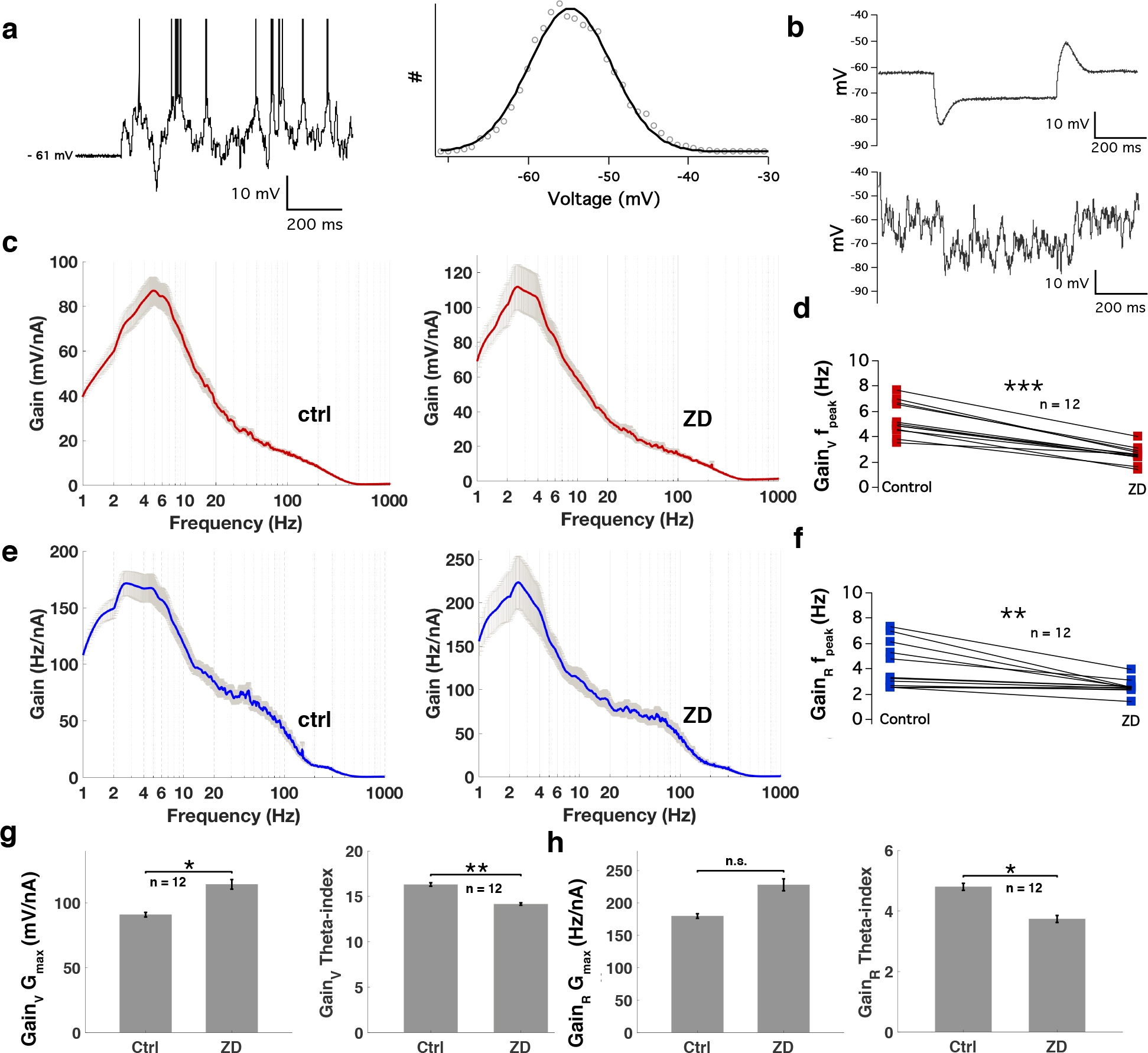
Mimicked in-vivo states and estimation of the signal gain in stellate cells. a. Voltage response of a stellate cell during mimicked in-vivo states (Left). Note spontaneous theta frequency firing and a detectable rhythmicity in voltage fluctuations. Membrane voltage histogram from the voltage trace in a, SD ~ 5 mV. (Right). b. Top: ~ 10 mV Steady state Voltage deflection from −500 pA current step before switching to the in-vivo regime, Bottom: Similar ~ 10 mV Steady state Voltage deflection from −1500 pA current step in the in-vivo regime. c. Gain_V_ in the control case (Left) and after blocking HCN channels with ZD7288 (Right). The plots represent average for 12 cells, error bars are s.e.m. d. Summary of the changes in the Gain_V_ f_peak_ for 12 cells in the presence of ZD7288. e,f. same as c,d for Gain_R_. g. Summary of the changes in the G_max_ (left) and theta-index (right) in the presence of ZD7288 for Gain_V_. h. Same as g for Gain_R_. d,f,g,h: *** p < 0.001, ** p < 0.01, * p < 0.05, n.s. non-significant, Student’s paired t-test.

We considered two types of responses, leading to two different signal gain waveforms. Gain_V_ (**Fig. 1c**), computed from the collective voltage trace, which mainly reflected the subthreshold frequency selectivity; and Gain_R_, (**Fig.1 e**), computed from a smoothly varying instantaneous firing rate (**Supplementary Figure 3)**, which reflected whether the observed subthreshold frequency selectivity can actually modulate the firing pattern of these cells.

The two estimates thus allowed us to assess the frequency dependent gain of subthreshold responses (Gain_V_) and firing rate (Gain_R_). Considering the contribution of HCN channels in maintaining the theta frequency resonance in stellate cells, we also re-estimated the signal gain after blocking HCN channels. To quantify the observed changes, we compared the frequency at which the peak was observed (f_peak_), the actual gain at peak (G_max_), and the relative gain in theta frequency (4-14 Hz) (theta-index); across the control and the test cases.

In Gain_V_, we observed a prominent peak in the theta frequency (f_peak_ = 5.35 ± 0.38 Hz, mean ± s.e.m., n=12), which shifted significantly to the lower frequencies (2.56 ± 0.19 Hz, n=12) after blocking HCN channels with ZD7288 (ZD, 25 μM) (**Fig. 1c,d**); although the firing rates in the control and under ZD were comparable (**Supplementary Figure 3**). A similar HCN sensitive theta frequency selectivity was also observed in the membrane voltage power spectrum, suggesting a reflection of the theta resonance (**Supplementary Figure 4**).

The actual gain (G_max_) was slightly higher in test (114.36 ± 12.87 mV/nA, n=12) than the control conditions (90.94 ± 6.13 mV/nA, n=12) (**Fig.1 g**), which can be expected considering the effects of higher input resistance after blocking HCN channels. Interestingly, however, the relative theta-frequency gain (theta-index), was significantly higher in control conditions (15.82 ± 0.18, n=12) than the test (13.19 ± 0.12, n=12) (**Fig.1 g**).

Interestingly, Gain_R_ too displayed a prominent peak in theta frequency (f_peak_ = 4.2 ± 0.15 Hz, n=12), which shifted significantly to lower frequencies after blocking HCN channels (2.56 ± 0.05 Hz, n=12) (**Fig. 1e,f**); but unlike Gain_V_, the increase in G_max_ for Gain_R_ was non-significant (**Fig. 1h**). Interestingly, the theta-index was still significantly higher for control (13.24 ± 0.26, n=12) than the test conditions (11.33 ± 0.25, n=12) (**Fig. 1h**).

These findings are in accordance with the previous studies which reported the influence of the subthreshold resonance mechanisms on the active properties^27^, and suggest the frequency dependent gain as a suitable index for mapping the relationship between the sensory modalities mediated excitation and the resulting firing rate response.

### Effect of input statistics on the signal gain

Next, considering that the input statistics can have a role in determining the actual signal gain^29,32^, we systematically perturbed the synaptic parameters of the *in-vivo* regime and studied changes in the frequency dependent gain.

To assess the effects of the excitatory drive on the signal gain, we first increased the strength of excitation by increasing the mean and the standard deviation (SD) of the excitatory conductance one at a time. We observed that although the firing rate increased linearly with the excitatory strength, the f_peak_, or the theta-index of Gain_V_ or Gain_R_ in control (**Fig. 2a,b**) or with ZD (**Supplementary Figure 5**) did not show any significant change.

**Figure 2.**
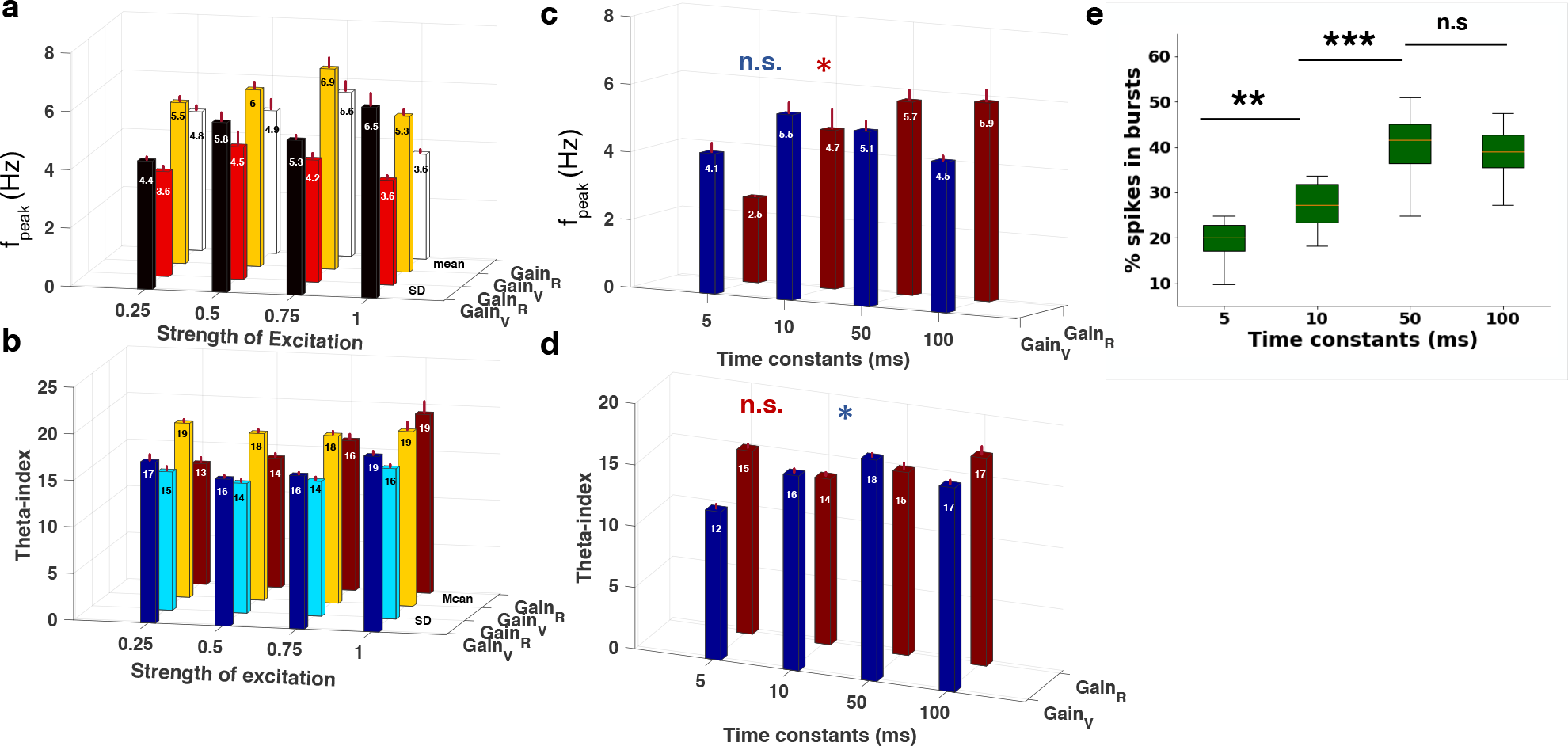
Effects of stimulus statistics on the signal gain a. Variations in the f_peak_ of gain_V_ and gain_R_ as a function of strength of excitation (normalized mean (n = 6) or SD (n = 9) of the excitatory conductance). b. Same as a for the theta-index of gain_V_ and gain_R_. c. Variations in the f_peak_ of gain_V_ and gain_R_ as a function of input kinetics (n = 7). d. Same as c for the theta-index. e. Percentage of spikes occurring in bursts as a function of the input kinetics. c,d: * p < 0.05, n.s. non-significant, Kruskal-Wallis test; e: *** p < 0.001, ** p < 0.01, n.s.: non significant, Student’s paired t-test.

The G_max_, however, showed a complex relationship with the excitatory strength, which can be explained by considering the voltage dependent influence of the HCN current (**Supplementary Figure 6**).

Next, in order to assess the effects of the synaptic kinetics, we repeated the gain measurements while gradually slowing down the kinetics of the excitatory as well as inhibitory conductances, by increasing the decay time constants of the synaptic conductances in steps of 5, 10, 50 and 100 ms.

Again, the gain_V_ f_peak_ was apparently unaffected by the input kinetics (**Fig. 2c**). However, the theta-index of Gain_V_ was observed to increase significantly with the input kinetics (**Fig. 2d**), possibly, owing to an increased summation of excitatory inputs in the theta frequency.

Next, analyzing the suprathreshold responses, we observed that the mean firing rate, while still being in theta frequency, decreased significantly as the time constants increased beyond 10 ms (**Supplementary Figure 7**), indicating significantly slower rate modulation. Surprisingly however, in spite of this decrease in the mean firing rate, the f_peak_ of Gain_R_, showed an interesting overall increase (**Fig. 2c**). One of the underlying factors causing this can be the higher frequency bursts, which, apart from increasing the gamma frequency power, when modulated in theta frequency, can give rise to an overall increase in power in theta frequency. To test this, we analyzed the percentage of spikes occurring as bursts (frequency > 50 Hz) for each group, and indeed observed that the bursts increased significantly as the synaptic time constants increased from 5 to 50 ms (**Fig. 2e**). The Gain_R_ G_max_, however, increased up to a certain range (50 ms), and decreased thereafter (**Supplementary Figure 7**).

These observations are in agreement with the previous findings^29^, whereby the peak frequency of the lower frequency resonance was observed to increase and the resonance strength was observed to decrease with slower input time constants, as a result of modulation of bursts. Our results thus confirm the role of bursts as a mechanism of active gain modulation.

### Theta modulation of synaptic inputs by HCN

Next, to probe further whether the amplified signal gain in theta frequency manifests in the modulation of synaptic inputs, we extracted the underlying theta oscillatory signal (**Fig. 3a**) from the collective voltage response by 4-15 Hz band pass filtering, and analyzed it further.

**Figure 3.**
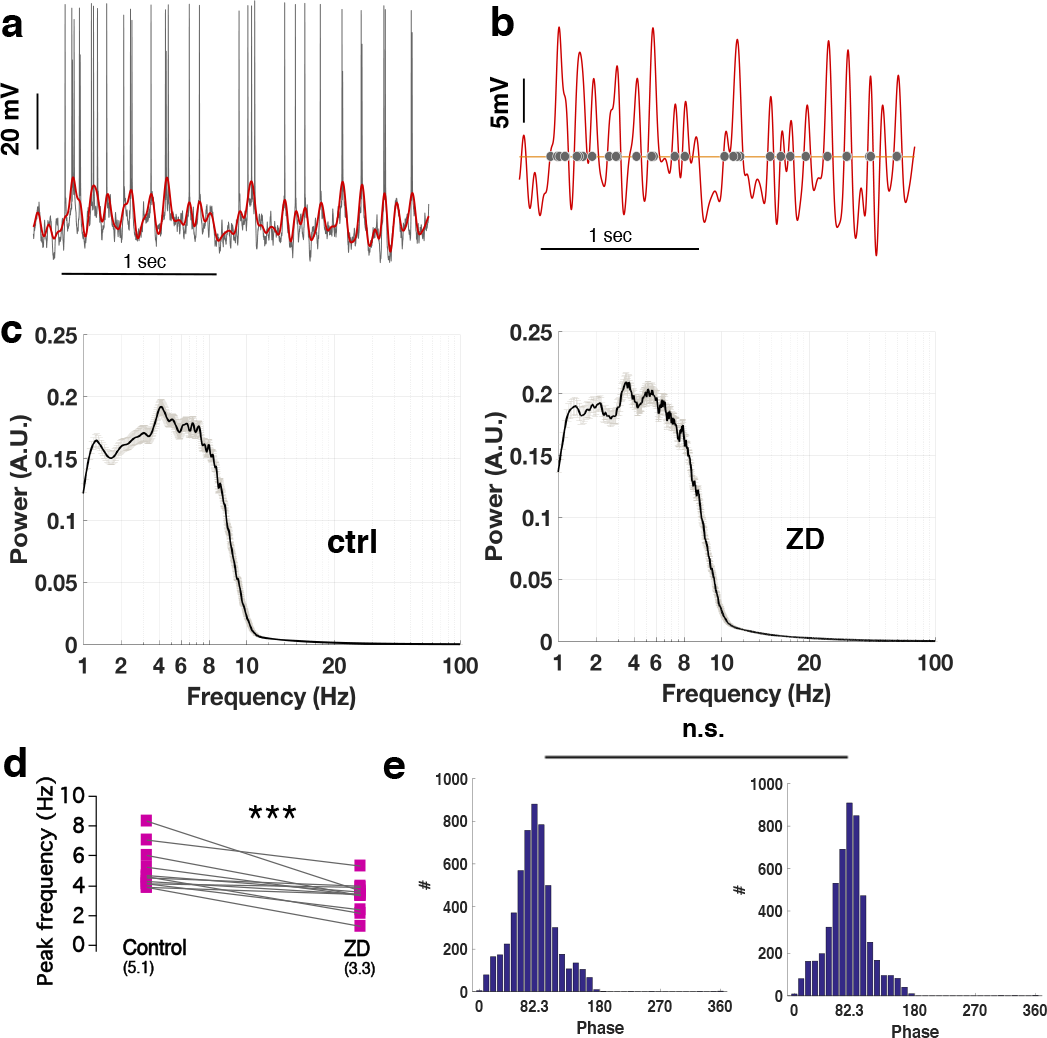
Theta frequency modulation of the synaptic inputs by HCN channels. a. 4-15 Hz band-pass filtered signal (red) overlaid on the actual voltage trace (black). b. The theta signal in a (red), and the corresponding spike timings (grey circles). c. Average power spectrum of the theta response (Left) and same after blocking HCN channels with ZD7288 (Right). The plots represent average for 12 cells, error bars are s.e.m. d. Summary of the changes in the peak frequency of the theta power spectrum in c, for 12 cells in the presence of ZD7288. e. Spike phase histograms for the pooled spike phases of 12 cells corresponding to their theta signals (see methods for the phase assignment). d: *** p < 0.001, Student’s paired t-test, e: n.s.: difference in the distributions non-significant, Kruskal-Wallis test.

Prominent subthreshold modulation in the theta range was indeed observed in this filtered signal, and on aligning the spike timings to it, the preference of spikes for the theta peaks was clear (**Fig. 3b**). For the default synaptic time constants, which were moderately slow, the peak in the power spectrum of this signal was obtained near 5 Hz on average (5.07 ± 0.12 Hz, n=12) (**Fig. 3c**). This peak frequency after blocking HCN channels was significantly reduced (3.3 ± 0.08 Hz, n=12) (**Fig. 3c,d**), indicating a salient theta frequency modulation of synaptic inputs by HCN channels. This peak frequency also showed a DV gradient for the sample size tested, which, interestingly, was flattened in the presence of ZD (**Supplementary Figure 8**).

Next, in order to determine the extent to which this theta oscillatory activity influences the spike timings, we analyzed the distribution of spike phases over the underlying theta (4-15) / delta (1.5-4 Hz) signal. We observed that most of the spikes fired at the near-peak ascending phase in control as well as with ZD (**Fig. 3e**), and the distributions of spike phases in both cases were almost identical.

Summarizing, for the given input parameters, HCN channels significantly contribute to the theta frequency modulation of the synaptic inputs. When the HCN induced resonance mechanisms are absent, the underlying subthreshold signal oscillates at a slower pace, but the spikes are still preferentially fired on the peaks of this subthreshold oscillatory signal.

### Theta modulation is sensitive to excitatory input kinetics

The results so far indicated that under *in-vivo* conditions, the synaptic inputs can be paced in a preferred frequency range by the neuron’s intrinsic resonance mechanisms, and even in the absence of such mechanisms, albeit in lower frequencies, a rhythmicity in the input structure can be present. These findings tally with the previous reports of persistent delta frequency rhythmicity in HCN1 knockout mice^33^.

Next, to assess the sensitivity of this theta modulation to the input statistics, we analyzed the underlying theta oscillatory signal while systematically varying the input parameters. For the control conditions, no significant change in the frequency of theta signal was observed on varying the SD and mean of the excitatory input as before, and also on varying synaptic kinetics within a certain range. But as the input time constants increased beyond this range, the frequency of modulation dropped significantly beyond the theta band (**Fig. 4a**).

**Figure 4.**
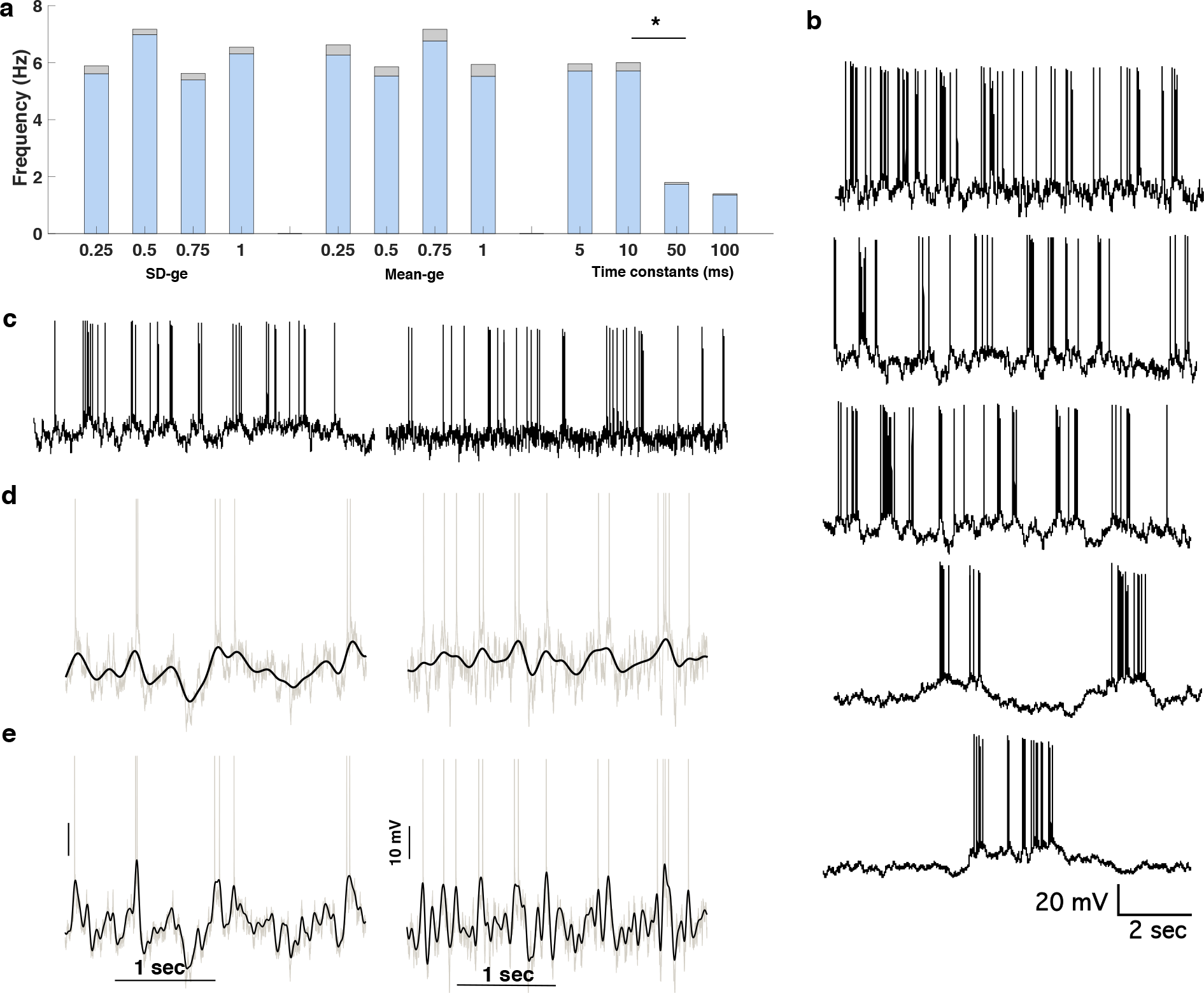
Theta modulation is sensitive to the excitatory input kinetics. a. Changes in the peak frequency of the theta power spectrum, as a function of the excitatory strength (normalized SD, mean of the excitatory conductance) and excitatory-inhibitory time constants. b. Voltage traces for the matched excitatory-inhibitory time constants varied from 50 ms, 100 ms, 200 ms, 500 ms, to 1000 ms top to bottom. c. Left: Voltage trace for the excitatory time constant = 100 ms, and inhibitory time constant = 10 ms; Right: Vice-a-versa. d. 3 Hz low-pass filtered ramp signals for the traces in c. e: 4-15 Hz band-pass filtered theta signals for the traces in c. a: * p < 0.05, Wilcoxon signed rank test, others: non-significant, Wilcoxon signed rank test & Kruskal-Wallis test.

Interestingly, however, with the slower input kinetics, the voltage response started exhibiting slow ramp-like motifs, which became more and more prominent on increasing the synaptic time constants (**Fig. 4b**). The increase in time constants was thus accompanied by a decrease in theta rhythm and emergence of slow ramps. It should be considered here that the synaptic kinetics so far were varied similarly for the excitatory and inhibitory inputs, thus making their individual contribution ambiguous. So next we made the excitation or inhibition selectively slower, while keeping the other at the default (10 ms; which is relatively faster), and observed that the time constants of excitation play a crucial role in determining the underlying rhythmic nature. When the inhibition was made faster, the visible ramps could still form; but that was not the case when the excitation was made faster against a slow inhibition (**Fig. 4c**); indicating that the emergence of ramps (**Fig. 4d**) requires a predominantly slow excitation. Also, interestingly, the underlying theta signal, became visibly faster as the excitation was switched to the fast mode (**Fig. 4e**), underlining the role of excitatory kinetics in controlling the pace of the fluctuating synaptic activity.

### Modulation of the excitation driven firing rate response

Next, in order to determine the extent to which the subthreshold theta modulation can alter the firing rate modulation, we did a detailed analysis of the temporal and spectral characteristics of the firing rate response. As stated previously, the firing rate increased with an increase in both, the mean and SD of the excitatory conductance, and the initial firing rates for the control and the ZD experiments were comparable (**Supplementary Figure 9**).

We observed however, that the percentage of spikes fired as bursts increased significantly with increase in the mean excitatory conductance in the control case, but not with ZD (**Fig. 5a**). Also, with increase in the SD of excitation, although the bursts increased significantly in both control as well as with ZD, the average number of bursts were significantly higher in the control case, for the entire middle range of the SD tested (**Fig. 5a**).

**Figure 5.**
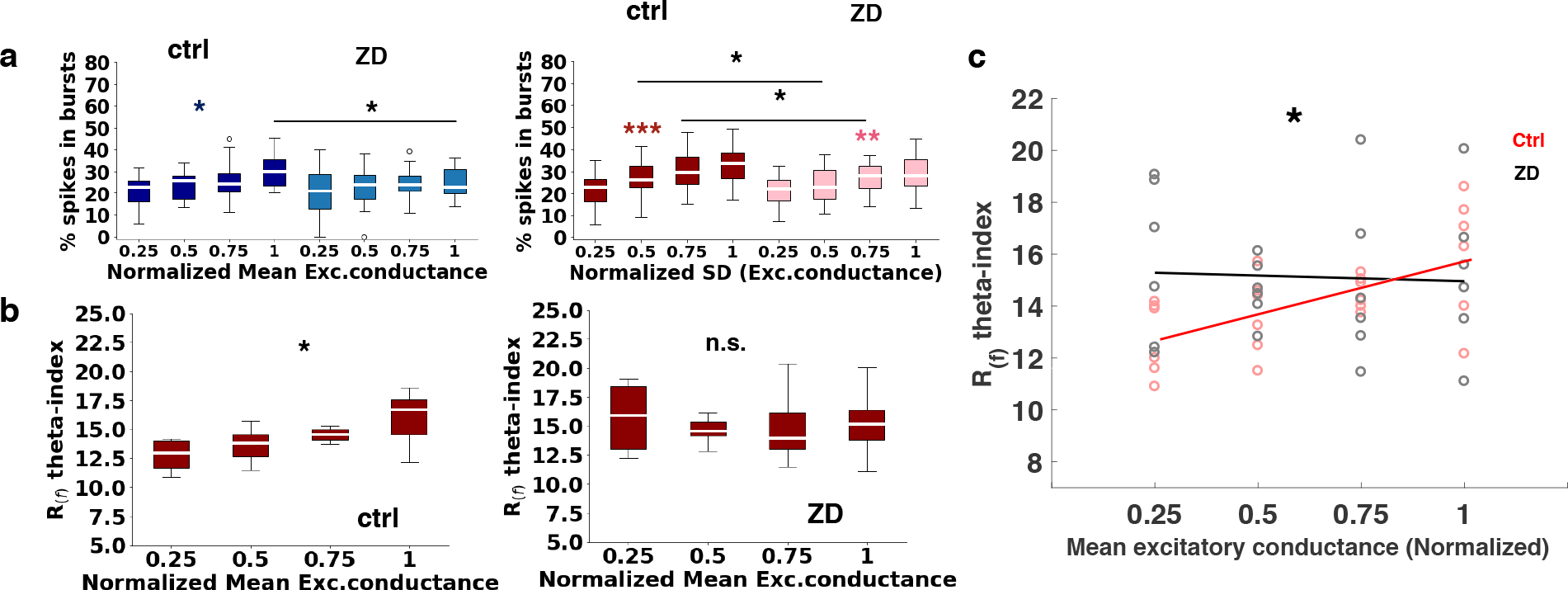
HCN controls the slope of theta specific firing rate modulation with excitation a. Percentage of spikes fired in bursts as a function of the mean and SD of the excitatory conductance (normalized) in control and with ZD (pooled data for n = 6 and 9 respectively). b. Theta-specific modulation of the firing rate response (theta-index of the rate response power spectrum), in control and with ZD. The overall increase was significant in the control case, but not with ZD. c. One-way analysis of covariance of the data in b shows a significant difference in slopes of excitation driven theta specific firing rate modulation between control and the test case. a: *** p < 0.001, ** p < 0.01, * p < 0.05, Kruskal-Wallis test for the pooled data, Student’s paired t-test for the individual experiments. b: * p < 0.05, n.s: non-significant, Kruskal-Wallis test, c. * p < 0.05, Analysis of covariance, F-test.

Also, to understand the relationship of these gamma frequency bursts with the underlying theta signal, we analyzed the distribution of the spikes over the theta/delta phases as before, and observed that, in all the cases, the neuronal firing was aligned to the peaks in the subthreshold signals, in control as well as with ZD (**Supplementary Figure 9**).

These findings clearly indicate that in the presence of HCN, there are more number of bursts in response to increasing excitatory drive, which are being modulated in the theta frequency, as opposed to lesser number of bursts with ZD, which are modulated in the slower delta frequencies. This confirms an overall stronger theta-gamma coupling in the presence of HCN channels. To study its implications, we analyzed the theta frequency modulation of the firing rate response, and observed that the theta-index of the power spectrum of the rate response increased significantly with the mean excitation in the control case, but not with ZD (**Fig. 5b**). Further analysis confirmed that the slope of this theta frequency modulation was significantly higher in the control case (**Fig. 5c**), which parallels the findings of the HCN1 knockout experiment^25^.

### Inhibition as an alternate resonance mechanism

In order to verify the mechanistic understandings gathered so far, we next constructed a computational model of MEC Stellate cell, and subjected it to similar *in-vivo* input regime. The electrophysiological signatures of stellate cells and the hallmarks of successfully operating *in-vivo* activity ^31^ could be tallied in the model (**Supplementary Figure 10**) enabling us to proceed with the signal gain measurements.

The Gain_V_ and Gain_R_ estimated from this model displayed similar theta frequency selectivity as observed experimentally (**Fig. 6a)**, thus confirming the sufficiency of proposed mechanisms. Next, for a better substantiation of the role of HCN channels, we decreased the HCN conductance in the model in two steps (1 (maximum), 0.5 and 0), and observed a likewise lower frequency shift of f_peak_, and a decrease in theta-index for both Gain_V_ and Gain_R_ (**Fig. 6a,c,d, Supplementary Figure 11**). The experimentally observed theta-modulation by HCN channels could also be verified in the computational model (**Supplementary Figure 10**).

**Figure 6.**
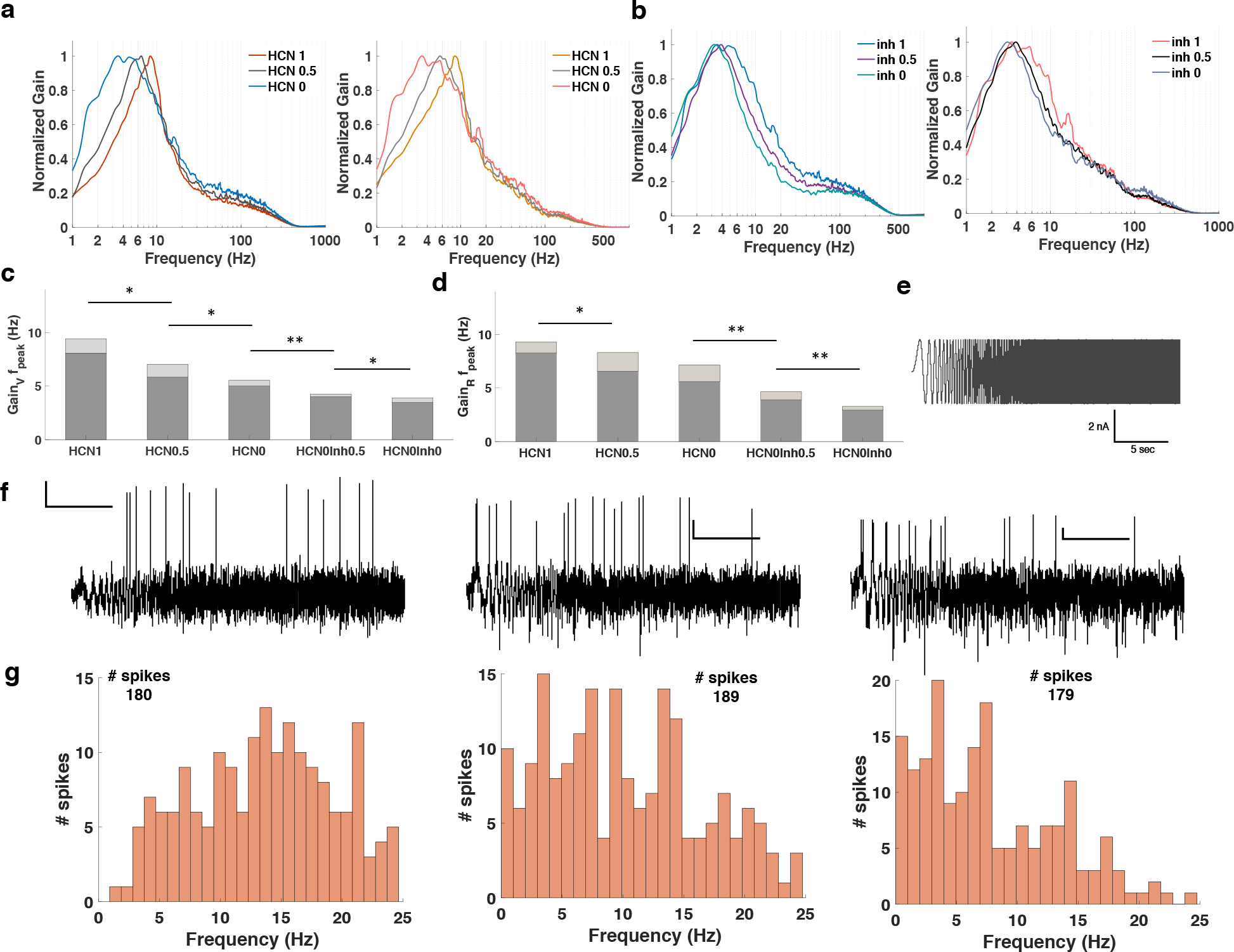
Inhibition controls the frequency selectivity in signal gain in the absence of HCN channels. a. Normalized Gain_V_ (Left) and Gain_R_ (Right) as a function of normalized HCN conductance. b. Normalized Gain_V_ (Left) and Gain_R_ (Right) as a function of normalized inhibitory conductance in 0 HCN condition. The plots in a,b represent average of 10 trials of 30 sec. each. c,d. Summary of the changes in f_peak_ for the traces in a and b. e. Chirp stimulus spanning 25 Hz in 25 sec. f. Voltage responses to stimulus in e injected under the in-vivo regime, for control (Left), no HCN (Middle), and no HCN-no Inhibition (Right) cases. Scale Bar: 5 sec., 20. mV in all panels. g. Histogram of spike times for total spikes fired in 10 trials as a function of stimulus frequency, for data in f. c: ** p < 0.01, * p < 0.05, Kruskal-Wallis test.

Next, to understand the factors regulating the remaining frequency selectivity in the absence of HCN channels, we decreased the strength of inhibition in two steps (1 (maximum), 0.5 and 0), and observed a further leftward shift in the frequency preference (**Fig. 6b,c,d**). Considering that a gradient of grid cell periodicity remains in the absence of HCN channels^25^, and a dorsoventral gradient of inhibition is present in MEC^30^, these observations should be noteworthy.

Hitherto, the analysis was carried out at the default synaptic time constants. It was possible that after removing HCN channels and inhibition, excitation can further affect the signal gain. To confirm this, we repeated the analysis using substantially slow excitation and observed that the remaining small frequency selectivity is noticeably affected by the slower excitation for Gain_V_ as well as for Gain_R_ (**Supplementary Figure 11**). The gamma frequency gain for gain_R_ was also visibly increased following the change in excitatory kinetics (**Supplementary Figure 11**), corroborating the experimental findings of such increase with slower input time constants (**Supplementary Figure 7**). Finally, to directly verify the effects of input selectivity analyzed so far, we tested the model with chirp stimuli (**Fig. 6e**), with increase in stimulus frequency from 0 to 25 Hz in 25 sec.

The roles of HCN channels were evident from the decreased firing on lower stimulus frequencies in the presence of HCN (**Fig. 6f,g** **left**), which increased after removal of the HCN channels (**Fig.6f,g** **middle**). Next, removing inhibition from the cell, further accentuated the firing response at lower frequencies (**Fig. 6f,g** **right**), thus suggesting the role of inhibition in contributing to weak resonance mechanisms in the stellate cells in the absence of HCN channels.

The results so far suggest that the HCN-based resonance mechanisms in stellate cells, besides modulating the voltage responses in theta frequency, also bias the preference of these voltage responses for the theta-frequencies in the stimulus. Collectively, HCN and inhibition thus reduce action potential firing both on and in the lower frequencies.

## Discussion

### Theta frequency resonance persists under *in-vivo* states

We observed that the HCN mediated resonance mechanisms are operational during the mimicked *in-vivo* states, despite a drastic drop in the input resistance and high noise levels, and can amplify the signal gain in the theta range.

Another major role attributed to HCN channels is their effect on the membrane resistance, which dictates the membrane time constant, and which in turn, controls the neuronal integrative properties^9^. However, under *in-vivo* states, where the membrane resistance drops by ~ 70-80%^31,34,35^, these reciprocal effects of HCN conductance have been questioned^36^. Our findings reveal that under *in-vivo* states, although HCN conductance leads to an overall reduction in the gain, HCN current can modulate the gain in a frequency dependent manner, by selectively increasing the gain over theta frequencies.

### Resonance as a dynamic gain control mechanism

Next, as we systematically perturbed the synaptic noise statistics, we observed that although the gain amplitude was sensitive to the input strength, its frequency selectivity was unaffected by such input scaling.

However, on making the synaptic kinetics slower, we observed a small but significant rightward shift in the otherwise unperturbed theta frequency peak, and an increase in the average gamma frequency gain. Both these findings corroborate the earlier reports of stronger higher frequency resonance when these frequencies in the input are weaker^29^.

This differential responsiveness of gain to the mean, SD and kinetics of stimulus can be a dynamic gain control mechanism, wherein the neuronal sensitivity to the input statistics is enhanced, enabling a state-dependent adaptation of their responses. Thus, rather than being dampened by noise, we observed the resonance to instead emerge through the latent noise statistics.

### Resonance manifests in the theta modulation of inputs

We observed a potent modulation of synaptic inputs in the theta frequency in the presence of HCN channels. Also, the spikes were phase locked to the peak of this signal, suggesting the possible influences on the firing rate modulation, which were later confirmed. In the presence of ZD, the frequency of this signal dropped to the delta range, but the spikes were observed to still occur selectively near the peak of the underlying delta signal.

Further, we observed that the frequency of this theta signal in control case was unaffected by the excitatory strength, but was sensitive to the excitatory input kinetics, which, when faster could contribute to the theta modulation, but when slower, mostly resulted in ramp-like motifs. Thus, our results bring to the fore the key targets: excitatory kinetics and synaptic integration, which when affected, can directly influence the synaptic ramp formation and theta oscillatory activity.

Summarizing, an oscillatory activity is present in the stellate cell voltage responses and is potent enough to modulate the spike timings, notwithstanding the heavy noise. The frequency of this modulation can vary from the dorsal to the ventral MEC, aligned to the HCN conductance, and can have a role in influencing grid-like responses of stellate cells as speculated earlier for the intrinsic oscillations.

### Stronger theta-gamma coupling steepens the slope of the firing rate modulation

A detailed analysis of the firing rate response revealed its susceptibility to the underlying subthreshold theta signal. The slope of theta frequency modulation of firing rate with the mean excitation was significantly higher in the control case, where the underlying subthreshold activity is theta-paced, than with ZD, where it oscillates in the delta frequency. Also, it became evident from the analysis of bursts, that there exists also a corresponding theta modulation of the gamma frequency bursts, which makes the overall theta-gamma coupling stronger in the presence of HCN channels. These findings parallel the similar higher slope of theta frequency modulation by running speed in control mice compared to the HCN1 knockout mice^25^, and tally the mechanistic implications of our findings.

### Possible effects on the grid spacing

We observed the theta-peak of the signal gain to be a direct reflection of the strength of HCN conductance, and found it to be robust to an array of input statistics. This resonance also reflected in the theta modulation of inputs and in steeper theta-gamma firing rate modulation with excitation, thus influencing the mapping between the excitatory inputs and the firing rate response.

Thus, there is room for speculating that the dorso-ventral gradient of grid spacing is actually a manifestation of the dorso-ventral distribution of HCN influenced differential resonance frequencies of the signal gain, which can reflect in the correspondingly faster theta-gamma modulation of the firing rate with the velocity mediated excitatory drive.

The reminiscent delta frequency selectivity in signal gain after blocking HCN channels parallels the HCN1 independent peri-threshold activity^33^ and the reminiscent gradient of grid periodicity observed in the HCN1 knockout mice^25^. This indicates a strong possibility of existence of alternate, HCN independent/co-dependent resonance mechanisms which can also be functional in the grid cell types devoid of HCN channels. Given that resonance is known to emerge through the interaction of two counteracting mechanisms^37^, we suspected a role of inhibition. Interestingly, we indeed observed a leftward shift of signal gain peak with decrease in the inhibitory conductance, indicating that the reported dorso-ventral inhibitory gradient^30^ can contribute to the similar gradient of frequency selectivity along the MEC DV axis, and can underlie the reminiscent gradient of grid periodicity observed in the HCN1 knockout mice. These proposed roles of inhibition are in accordance with those already known in the case of cortical circuits^38^.

The combined effects of the excitatory-inhibitory dynamics discussed above, can dictate different symmetry breaking operations along the dorso-ventral MEC axis, which can be further fine-tuned by HCN channels, thus setting the grid scale.

### Other implications

The robustness of stellate cell resonance to noise level can be an error-correction mechanism, necessary for maintaining the accuracy of the spatial coding^39^. A possibility thus also remains that the strong HCN dependent resonance mechanisms in stellate cells, which are also reflected in the firing rate modulation, can influence the network level input statistics^40^, thus influencing input statistics based resonance in other cell types (in MEC as well as stellate cell’s downstream targets in the hippocampus). Stellate cells in that case can indirectly serve their long believed role of pacemakers.

## Online Methods

### Brain slice preparation

The guidelines of ‘Animal Ethics and Welfare Committee’ of the Indian Institute of Science, Bangalore, India were followed in all the experiments. 5-8 week old male SD rats (weighing ~130 - 250 g) were used in the study.

Rats were anesthetized by intra-peritoneal injection of a combination of Ketamine and Xylazine (dose calculated according to the animal weight). After onset of deep anaesthesia, as determined by cessation of toe-pinch reflex, rats were transcardially perfused with ice-cold cutting solution (see Solution composition) continuously bubbled with carbogen (a mixture of 95% O_2_ and 5% CO_2_ gas), and then decapitated.

During dissection, the skull was carefully separated through its midline and the brain was scooped out using a blunt spatula from the side of the cerebellum and was immediately transferred to sACSF (see Solution composition) pre-chilled to 0-4 °C and continuously bubbled with carbogen. This part of the dissection was usually completed within a minute.

Parasagittal hippocampal slices (350 μm) were then obtained according to the guidelines provided by previous reports^41^, while submerged in oxygenated ice-cold cutting solution, using Leica VT1000S vibratome. Brain slices were then transferred to the incubation solution (see Solution composition) in an incubation chamber, maintained at 34 °C using a water-bath. Slices were placed according to the hemisphere maintaining the order in which they were obtained. The incubation solution was pre-bubbled with carbogen which was continued throughout the incubation period.

After 15 minutes of incubation, the incubation chamber was transferred to room temperature (~ 24 °C), continuing the carbogen bubbling. Electrophysiology recordings were started after 1-hour recovery period from the end of this incubation.

### Solution composition

The following solutions were used for the electrophysiology experiments. (all salts from Sigma Aldrich).

### Sucrose based Artificial Cerebrospinal Fluid (sACSF)

The solution used for acute slicing procedure contained (in mM): Sucrose (230), KCl (2.5), MgCl_2_.6H_2_O (7), CaCl_2_.2H_2_O (0.5), NaHCO_3_ (25), NaH_2_PO_4_ (1.25), Glucose (10), Sodium Pyruvate (3). pH 7.4, osmolarity ~ 290 - 330 milliosmoles.

### Incubation Solution

The slices were incubated in an incubation solution containing (in mM): NaCl (125), KCl (3), MgCl_2_.6H_2_O (1.5), CaCl_2_.2H_2_O (2), NaHCO_3_ (25), NaH_2_PO_4_ (1.25), Glucose (10), Sodium Pyruvate (1). pH 7.4, osmolarity ~ 290 - 310 milliosmoles.

### Artificial CerebroSpinal Fluid (nACSF)

For electrophysiology recordings, the slices were transferred to the nACSF containing (in mM): NaCl (125), KCl (3), MgCl_2_.6H_2_O (1), CaCl_2_.2H_2_O (2), NaHCO_3_ (25), NaH_2_PO_4_ (1.25), and Glucose (10); pH 7.4; osmolarity ~ 290 - 310 milliosmoles.

### Internal Solution

The patch pipettes were back filled with an internal solution consisting of (in mM):

Potassium gluconate (120), KCl (20), HEPES (10), Disodium phosphocreatine (10), MgATP (5), Na_2_GTP (0.5), EGTA (0.2). pH 7.4; Osmolarity: ~ 290 - 310 milliosmoles.

### Pharmacological agents

During experiments, these were perfused with nACSF when required.

Synaptic blockers:

CNQX 10 µM, (+)bicuculline 10 µM, picrotoxin 10 µM, D,L-AP5 50 µM, CGP55845 5 µM (Abcam). HCN blocker: ZD7288 25 µM (Tocris Bioscience).

### Data acquisition and analyses

Experiments were performed in nACSF at 34 °C; temperature maintained with the help of a bath chamber heater connected to an automatic temperature controller (Harvard Apparatus, USA). The patch pipettes (1.5 mm O.D., 0.86 mm I.D.) were prepared using P-97 puller (Sutter Instruments, USA) and fire polished to achieve a final access resistance of 3.5 - 5 MΩ. These were then back filled for stellate cell recordings, with the internal solution.

Medial Entorhinal cortex layer II stellate cells were identified using infrared Dodt gradient contrast video microscopy (Olympus BX51WI). These putative stellate cells were then patched in whole-cell current-clamp configuration to record intrinsic electrophysiological properties and their modulation. Somatic whole cell recordings in fast current-clamp mode were performed using DAGAN-BVC 700A amplifier, (Dagan Corporation, USA), Digidata 1440A Digitizer (Molecular Devices, USA), and WinWCP software package (Strathclyde Electrophysiology Software, UK). The signals were filtered at 5 kHz and stored on disk at > 20 kHz sampling frequency. Series resistance (< 25 MΩ) was fully compensated in the current clamp mode. Series resistance was monitored and compensated online using the bridge-balance circuit of the amplifier.

The following parameters i.e., resting membrane potential (RMP), input resistance (R_in_) and series resistance (R_s_) were regularly monitored during the electrophysiological recordings. Recordings in which the series resistance did not change > 5 MΩ throughout the duration of the experiment were included for analysis. The stellate cells had their break-in RMP of ~ −62 - −65 mV. Unless otherwise stated, experiments were performed at the initial resting membrane potential of the cell. Cells showing RMP depolarized than −60 mV during the recordings were discarded from the analyses.

The system for the dynamic clamp experiments consisted of the Real-time Experiment Interface (RTXI) (version 2.0-rc3, http://www.rtxi.org/) installed on a personal computer, Dagan BVC 700A amplifier and PCI-6221 multi-function data acquisition (DAQ) board (National Instruments, USA), with a BNC-2090A coaxial cable interface.

Dynamic clamp signals were filtered at 5 kHz and sampled at 10 kHz.

### Mimicking in-vivo states

After recording the basic neuronal parameters in current clamp, the recording mode was switched to the dynamic clamp.

The fluctuating synaptic conductance (Gfluct) module in the dynamic clamp (adapted from Destexhe *et. al.*, 2001^31^ and Gfluct.mod file (ModelDB Accession: 8115)), was used to generate in-vivo like background synaptic activity.

### The Default Gfluct parameters

The mean excitatory conductance: g_e0 = 0.03 μS, The mean inhibitory conductance: g_i0 = 0.06 μS, Standard deviation of the excitatory conductance: std_e = 0.01 μS, Standard deviation of the inhibitory conductance: std_i = 0.02 μS, Time constant for the excitatory conductances: Tau_e = 10 ms, Time constant for the inhibitory conductances: Tau_i = 10 ms, Excitatory reversal potential: e_rev = 0 mV, Inhibitory reversal potential: i_rev = −75 mV. The voltage responses and the applied current were recorded for 8-10 trials of ~10 sec. each. This duration was increased when the slower synaptic conductances were used, to account for the slower firing rate and to sample approximately the same number of spikes.

The changes in basic electrophysiological properties were monitored periodically. The changes in series resistance were monitored and adjusted for in between the trials.

### Experiments to test the role of input statistics

In a set of experiments designed to test the effects of varying mean and SD of excitation, the SD of the excitatory conductance was varied from 0.002 μS to 0.02 μS in steps of ~0.005 μS, while holding the mean unchanged at the default value. Then, the mean of the excitatory conductance was varied from 0.02 μS to 0.04 μS in steps of 0.005 μS, while holding the SD unchanged at the default value.

To test the role of synaptic kinetics, the time constants of excitatory and inhibitory synapses were changed from 3-5 ms, 10 ms, 50 ms, 100 ms and 200 ms; keeping rest of the parameters default. These values are selected considering the time constants of the synapses observed in-vivo, and those used previously for the stellate cells ^26,31^.

In a small subset of experiments, extreme values of synaptic time constants such as ~ 0 or 500 - 1000 ms were used to confirm the mechanistic interpretations.

### Experiments to test the role of I_h_

To test the role of HCN current (I_h_), ZD7288 was used as HCN channel blocker.

After recording all the control traces, 25 μM of ZD7288 was bath perfused. The test recordings were started ~ 10 min. after starting this perfusion, and after ensuring a complete HCN block via the electrophysiological signatures.

### Data Analyses

Acquired electrophysiological data were analyzed using Igor Pro 6 (Wave Matrix, USA), Matlab (The MathWorks, Inc.), Python programming language (Python Software Foundation), and Clampfit (Molecular Devices, USA).

The input resistance was measured at the steady state of voltage deflection.

### Estimation of signal gain

The neuronal voltage response was analysed in two different ways to estimate the signal gain.

In the first case, the collective voltage trace was analysed in the time and frequency domain. For the further analysis, however, a vector of smoothly varying instantaneous firing rate as a function of time was constructed from the actual voltage response (**Supplementary Figure 3**).

To calculate the signal gain, a ratio of the Fourier transform of the response and the Fourier transform of the stimulus (applied current) was calculated (Analysis done in Matlab). The gain function thus obtained was filtered using a Gaussian window and smoothened using a moving average. The window width was adjusted so as to achieve a noise free trace, without compromising on the signal information. On average, a window of 20 X-axis data points was used, corresponding to ~ 1.5 Hz.

### Analysis of underlying theta oscillations and slow synaptic ramps

To compare the changes selective to the theta frequency range, the output voltage traces were band-pass filtered using a 4-15 Hz FIR filter. Similarly, to analyze the slowly varying ramps, the output voltage traces were low-pass filtered using a 3 Hz FIR filter.

### Estimation of spike phases

For estimating the distribution of spike phases, the theta and ramp signals were used to determine their peak times (corresponding to the positive peak), trough times (corresponding to the negative peak), and zero-times (all time points where the signal crosses zero: The mean of the mean subtracted, normalized signal).

Thus, three zero-points constituted one oscillatory cycle. Phases from 0-360 were then assigned to each cycle. A phase vector corresponding to the original voltage trace could be thus obtained, which allowed to compute the phase corresponding to each particular spike.

The mean, median, mode and variance of spike phases were then computed from a histogram of pulled spike phase values.

### Analysis of bursts

To study the percentage of spikes occurring as bursts, spikes occurring in 50-100 Hz bursts were considered as a single spike, and spike time of only the first spike in the burst was considered. The number of spikes and the firing rate was then recalculated.

### Computational model

A single compartment model of MEC layer-II stellate cell was built in NEURON simulation environment (Hines and Carnevale, 1997)^42^, with parameters adjusted to match the observed electrophysiological responses of the Stellate cells.

### Fluctuating conductances

The in-vivo like fluctuating synapses were modelled according to the Gfuct.mod input regime already discussed under “Mimicking in-vivo states”.

### Voltage gated Na^+^ and K^+^ channels

Adapted from a computational model by Destexhe et. al., 2001^31^.

ModelDB Accession:8115

### HCN channels

Adapted from Christoph Schmidt-Hieber and Häusser, 2013^22^.

ModelDB Accession:150239

### Persistent Na^+^ channels

By Uebachs et al., 2010^43^.

Assumes a non-inactivating conductance.

ModelDB Accession:125152

### K_AHP_ channels

Adapted from Migliore, Cook, Jaffe, Turner, & Johnston, 1995^44^.

ModelDB Accession: 3263.

### L-type Ca^++^ channels

Voltage gated high threshold activated calcium channel, adapted from Migliore, Cannia, Lytton, Markram, & Hines, 2006^45^.

ModelDB Accession: 64229

### Calcium handling mechanism

Models the decay of the internal calcium concentration due to calcium currents and pumps.

Adapted from Destexhe, Contreras, Steriade, Sejnowski, & Huguenard, 1996^46^.

ModelDB Accession: 17663

### Statistical Analysis

The tests used for hypothesis testing and calculating the p-values are described with the individual results.

Typically, student’s paired t-test, Kruskal-Wallis test, Analysis of covariance (ANCOVA), and F-statistics were used as appropriate.

The statistical analysis was carried out using Matlab, MS-Excel and the Python programming language.

